# A paper based microfluidic platform combining LAMP-CRISPR/Cas12a for fluorometric detection of nucleic acids

**DOI:** 10.1101/2023.03.02.530841

**Authors:** Anindita Sen, Calum Morris, Aashish Priye, Murray Broom

## Abstract

Nucleic acid isothermal amplification methods are advantageous for point-of-care (POC) diagnostics because of their precision, sensitivity, and low power requirements. Loop-mediated isothermal amplification (LAMP) is an established method, renowned for its nucleic acid amplification efficiency and robust amplification of semi purified target nucleic acids. However, LAMP may be prone to non-specific amplification causing false positives and therefore fails to replace PCR as the gold standard method in clinical testing. We show that LAMP combined with clustered regularly interspaced short palindromic repeats (CRISPR) technology is an effective alternative for overcoming the limitations of LAMP alone. Nucleic acids are first isothermally pre-amplified to enrich for targets, then specific amplification detection signals are generated by sequences of RNA guided recognition of amplicons. We are the first to demonstrate a paper based microfluidic system for detecting pathogen nucleic acids in samples by combining the power of LAMP and CRISPR technology. We show that although LAMP may produce non-specific amplification, the possibility of detecting a false positive can be eliminated by combining LAMP with CRISPR based detection. We demonstrate that a paper based microfluidic platform has the potential to compete with the conventional Nucleic acid testing (NAT) technologies not only in terms of robustness but also in terms of cost and complexity.

## Introduction

LAMP has emerged as a popular isothermal amplification technique for pathogenic nucleic acid detection since its introduction in the 2000.[1], [2] Owing to its lack of need for thermal cycling and superior detection capabilities with high sensitivity and specificity, LAMP has obvious advantages over more widely used amplification methods such as PCR.[3] It is faster, less expensive, less complex in terms of instrumentation and less vulnerable to the presence of extraneous nucleic acids than PCR.[4] Whilst claiming to offer superior specificity compared to PCR owing to the use of four-six target specific primers binding to six to eight regions (compared to only two primers in PCR), LAMP is unsurprisingly prone to non-specific amplification due to the same reasons[5]. False positive detection in LAMP creates a general setback particularly for diagnostic use and there is a need for technologies that can combine the cost efficiency and simplistic nature of LAMP with the diagnostic accuracy of PCR.

The use of nonspecific indicators to detect LAMP amplicons fail to differentiate specific target based amplification from primer dimer amplification [6],[7]. To make LAMP suitable for diagnostic applications it is necessary to boost sensitivity and specificity through additional readouts.[8], [9],[10] Thus, strategies coupling nucleic acid amplification technologies (NAAT) with CRISPR/Cas technologies such as DETECTOR[11], SHERLOCK[12] and HOLMESv2[13] to boost LAMP’s efficacy are emerging as the next generation of robust molecular diagnostics. These CRISPR/Cas systems operate based on a guide RNA (gRNA) dependent activation of the CRISPR-associated (Cas) enzyme that unleashes strong *trans*-cleavage activity cleaving single stranded nucleic acids. This biochemical event can be transduced into strong signals by using quenched fluorescence probes. So far, this strategy has been used by various studies and the superiority of LAMP-CRISPR technology has been demonstrated restricted to a laboratory setting relying heavily on tube based reactions.[14]–[17] However, it is imperative to move towards diagnostic platforms that are suitable for translating to resource limited settings.

Paper offers an attractive substrate for developing nucleic acid diagnostics. Paper is biodegradable, low-cost, hydrophilic and offers superior biocompatibility making it a promising substrate for successful loading of LAMP-CRISPR chemistries. Paper is inherently flexible and a suitable medium for fluid flow through capillary action not requiring any external force offering appealing possibilities in microfluidics.[18] Although there are studies that report LAMP on paper[19]–[25], to date no LAMP-CRISPR detection on paper substrate has been reported. Considering the temperature incompatibilities between LAMP and CRISPR, it was difficult to incorporate both into a combined system until reports like DropCRISPR[26] and SlipChip[27] platforms, that have used digital droplet based analysis for quantitative detection of nucleic acids. These approaches are limited by complexity and the use of precision instrumentation in most cases. However, to make nucleic acid testing (NAT) technologies accessible to resource limited settings and in developing countries they must be simple and cost-effective. This can be achieved through paper-based platforms.

Here we describe a paper-based LAMP-CRISPR-Cas12a assay platform to achieve qualitative fluorescent visual detection of amplified nucleic acids (Figure 1). We explicitly demonstrate the issue of spurious signals arising from LAMP using *E. coli* DNA as target and show that by harnessing the power of CRISPR-Cas12a system we overcome the detection of false positives. Following promising outcomes on paper we demonstrate specific detection of *E. coli* DNA in soil samples processed and extracted using S-TECH, a new technology for sample preparation at point-of-care. We report a sensitive, specific, and faster detection with LAMP-CRISPR on paper compared to LAMP alone in tubes. Our work provides a first proof-of-concept of a paper-based nucleic acid diagnostic platform that combines the simplicity of LAMP amplification with high sensitivity and selectivity of CRISPR detection systems.

**Figure 1.**
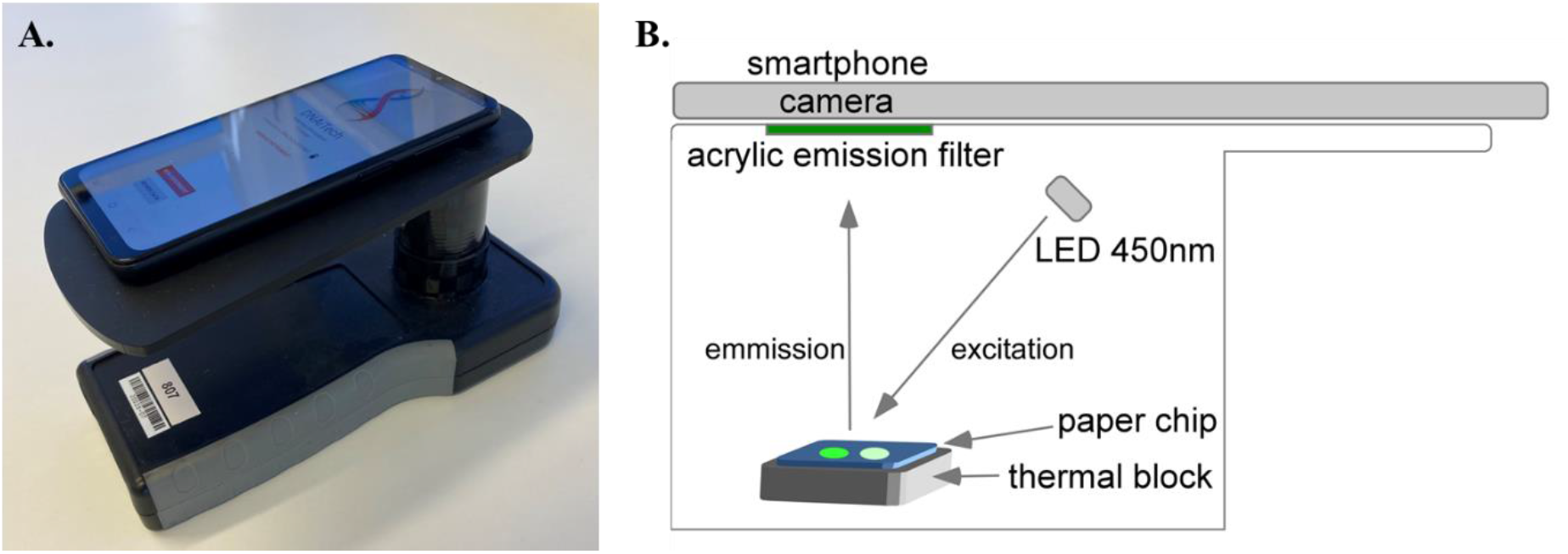
**A**. Image of DNAiTECH gen2 portable LAMP device. **B**. Schematic of the LAMP-CRISPR detection device prototype built in-house by DNAiTECH for carrying out LAMP reactions and CRISPR detection on paper.

**Scheme 1.**
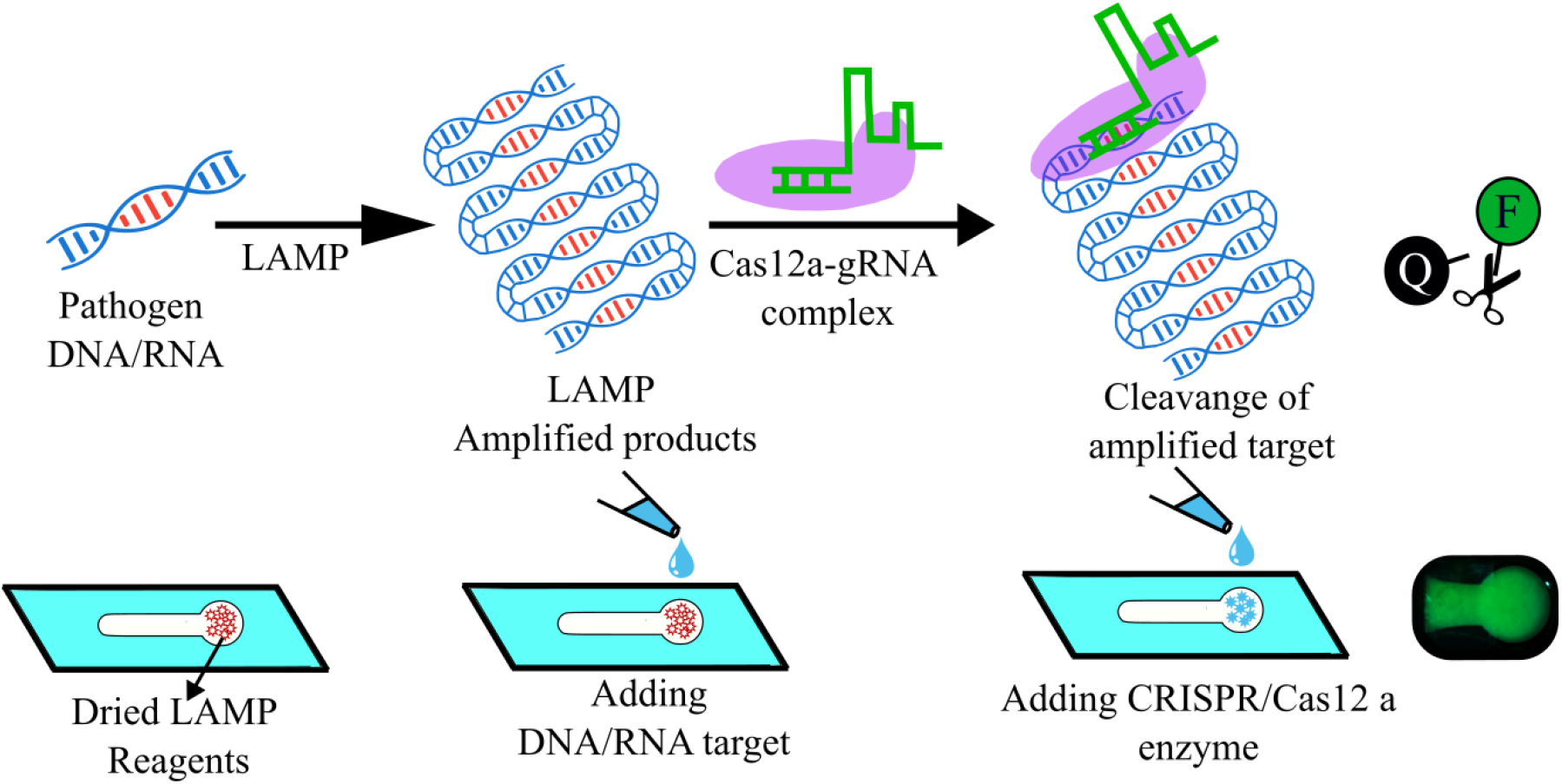
Schematic of the paper-based LAMP-CRISPR detection workflow. Paper strips containing dried LAMP reagents serve as simple and cost-effective nucleic acid testing assays. First, a sample containing template DNA/RNA is introduced to the testing zone followed by selective isothermal amplification of targeted sequence via LAMP. Second, subsequent addition of cas12 a enzyme to the testing zone activates ribonuclease activity to selectively cleave the amplified target to release the fluorescent reporter molecules. The bright fluorescent signals on paper strips can be visually read on smartphone for easy discrimination of positive and negative samples. By combining the programmability of CRISPR with the versatility of paper-based LAMP amplification, the sensitivity and selectivity of nucleic acid testing platforms can be greatly improved.

## Materials and Methods

### DNA Samples and *E*. *coli* cells

*Escherichia coli* DNA and cells was kindly provided by the HEV (Hub for Extracellular Vescicle Investigations), University of Auckland, New Zealand.

### LAMP assay

For tube-based LAMP assays, reactions were carried out in a total of 25 μL containing 1 X Isothermal Amplification Buffer (New England Biolabs), 5 mM MgSO_4_, 1.4 mM dNTP solution (New England Biolabs), 1.6 μM FIP/BIP, 0.2 μM F3/B3, 0.4 μM LF/LB, 2.4X EvaGreen (Biotium), 416 U/mL Bst 2.0 Warmstart DNA Polymerase (New England Biolabs) and made up to 20 μL with nuclease free water (IDT). For the reaction, 5 μL of template or nuclease free water was first added to a dome capped PCR tube and 20 μL of the LAMP reaction added and mixed. LAMP assays were overlaid with 60 μL mineral oil and run at 65°C with a 5-minute delay and 3 images per minute on the DNAiTECH portable isothermal amplification instrument (www.dnaitech.com).

*Escherichia coli malB* primers used in this study are shown in Table 1 and were synthesised by Integrated DNA Technologies (IDT, https://www.idtdna.com/).

**Table 1.**
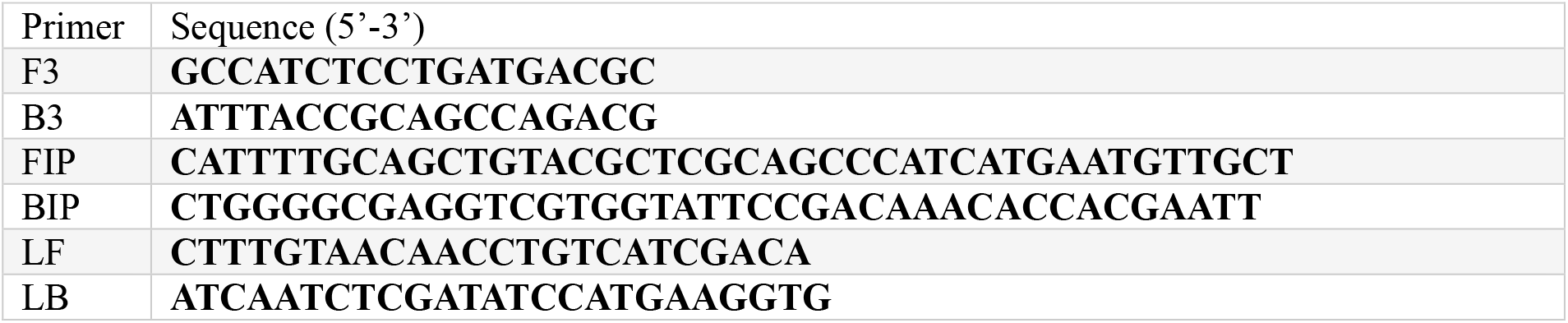
LAMP Primers and sequences used in this study [source *[28]*].

### *E*. *coli* crRNA design

The *E. coli malB* crRNA was designed using the inbuilt CRISPR design tool in Benchling (https://www.benchling.com/) to identify suitable PAM sites between the F2 component of the FIP and the B2 component of the BIP and synthesised by IDT. The complete crRNA sequence is shown in Table 2.

**Table 2.**
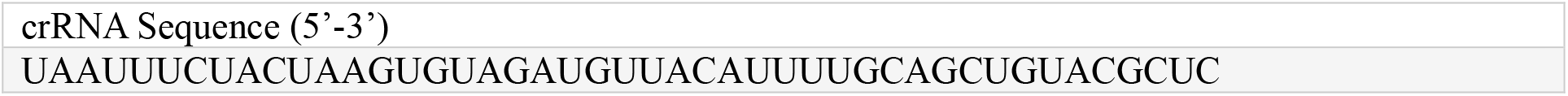
crRNA sequence used in this study.

### CRISPR assay

100 μM EnGen® Lba Cas12a (New England Biolabs) was diluted as needed in the supplied Cas12a diluent. For tube assays CRISPR mixes were made up to 10 μL containing 0.2 μM Cas12a, 1.2 μM crRNA, 2x NEB r2.1 Buffer (New England Biolabs), 5 μM fluorescent ssDNA probe (6-FAM/TTATT/BHQ1), and volume adjusted to 10 μL with nuclease free water. This was combined with 10 μL of LAMP product and incubated at 37°C in a DNAiTECH Gen2 portable real-time isothermal amplification instrument (www.dnaitech.com) with 3 images per minute and no delay.

### Single tube LAMP-CRISPR assay

For single tube LAMP-CRISPR assays, 8 μL of reaction mix and 2μL of template was added to the PCR tube and overlaid with 60 μL of mineral oil. 10 μL of CRISPR mixture (as described in CRISPR assay section) was placed within the lid of the PCR tube to separate it from the high temperature LAMP reaction. These reactions were placed at 65°C in the DNAiTECH instrument for 20 minutes to pre-amplify the target. The tubes are cooled to RT before adding the CRISPR mix. The separate LAMP and CRISPR reaction mix were combined by a short pulse in a centrifuge. The combined reactions were then incubated at 37°C with 3 images per minute.

### Specificity and sensitivity of LAMP

The sensitivity of the LAMP assay was assessed in tubes and LAMP-CRISPR on microfluidic chips. These were done using 10-fold serial dilutions of *E. coli* genomic DNA in IDTE (pH 8.0, IDT) including 40,000 genomic equivalents (GE), 4,000 GE, 400 GE, and 40 GE.

### Fabrication of paper chip

Whatman 113 and Whatman 1 were chosen as the paper substrates. Paper chips were assembled using laser cut tape and PET supports. 50 W CO_2_ Flux Beambox Pro was used to laser cut the components and Beam studio was used to design the respective cut-outs. Typically, the paper was secured on an adhesive tape before placing the PET cover with a hole for sample input on top. The fabricated paper chip was then pressed adequately by hands for securing the PET cover and the adhesive layer to avoid any leakage around the paper disc after sample flow.

### Drying LAMP reagents on paper

The LAMP cocktail was premade with the following components. dNTPs, LAMP primers same as for tube assays, Bst warm start (2.6 U/ul final concentration), 10% trehalose, 1 x LAMP buffer same as for tube assays. No magnesium sulphate was added to the drying mix. 10 μL of the mix was added to the paper discs which were then freeze dried using Labconco FreeZone 2.5 L benchtop freeze dryer. A 50 μL LAMP sample mix consisting of 5 mM MgSO_4,_ 1 x isothermal buffer, 10 μL target DNA and nuclease free water was used rehydrate the paper substrate prior to LAMP reaction.

### LAMP-CRISPR box and fluorescence detection

Our LAMP CRISPR chip reactions were carried out using a system similar to that published by Priye et al.[5] Heating of the chip to 65°C was achieved using 12 volts supplied to two 10 W resistors (Digi-key; Part # 3.3 W-10-ND) in series within an aluminium block. The Arduino’s digital pin was used to control the temperature of the block through an N channel MOSFET (MOSFET N-CH 60V 55A TO-220; Digi-key; Part # 497-6742-5-ND). A temperature sensor (Digi-key; Part # AD22100STZ-ND) was used to monitor the heater temperature. We use EMITTER 5MM 450 nm FLAT LENS TO39 blue light (Digi-Key Part Number 1125-1302-ND) illumination from above for excitation source and measured the fluorescence emission at >500 nm on the smartphone camera using a green acrylic emission filter. The box and all components were assembled from laser cut black acrylic sheets (3 mm).

### Soil sample preparation

Soil samples plus and minus *e. coli* cells were extracted using the S-TECH device (www.dnaitech.com). The extracted DNA was measured using the Qubit (Invitrogen Qubit 3 Fluorometer) and used for LAMP reactions in tubes and LAMP-CRISPR reactions on chips.

## Results and Discussion

All our tube-based LAMP-CRISPR assays were performed in our DANiTECH gen2 system (www.dnaitech.com). Briefly, our NAT device consists of low powered resistive heating elements that directly heats the paper substrates or liquid aliquots in tubes to maintain desired temperature setpoints to sustain LAMP and CRISPR reactions. A ring of blue LED lights illuminates the paper substrate to excite the fluorescent reporter molecules generated by LAMP or LAMP-CRISPR amplification systems. A smartphone camera in conjunction with a data analysis app (DNAiTECH real-time analysis version 1.3.4), records and images the post amplification signals generated from liquid aliquots in tubes to capture and quantify fluorescent emission signals (**Figure 1A**). A LAMP-CRISPR device prototype with a standard smartphone camera was used to image the fluorescent signals arising on paper from CRISPR detection of LAMP product to score the assay as a positive or negative (**Figure 1B**).

First, we demonstrate LAMP amplification and detection of *E. coli* DNA in liquid aliquot tubes. While the LAMP assay for *E. coli* is rapid, random nonspecific amplification in negative control samples may often occur. In our repeated LAMP reactions, we find that three out of eight and two out of six negative controls exhibited delayed false positive signals (**Figure 2 A, B**). We aimed to discriminate between contamination and non-specific amplification by employing CRISPR detection technology. Using CRISPR as the readout for the LAMP reaction, false positives were eliminated (**Figure 2 C**). **Figure 2 D** shows that the normalized intensity derived from the image pixel analysis of the positive and negative samples on paper. CRISPR/Cas12a confers detection specificity through its gRNA when the PAM sequence targets regions within the amplicon outside the lamp primer regions. Upon identification, the Cas12a endonuclease is activated and makes a ssDNA cut upstream/downstream of the PAM site on the amplicon. Thereafter, it unleashes indiscriminate *trans*-cleavage resulting in unquenched fluorescence that signal the presence of amplicons in the sample[29]. Hence, any non-specific amplification during LAMP is undetected during CRISPR which only specifically detects target induced amplification. Lack of fluorescence in the negative control when combining LAMP with CRISPR based detection ruled out the possibility of contamination. This demonstrates that although LAMP may produce non-specific amplification, the possibility of that resulting in a false positive can be eliminated by harnessing the power of specific detection by CRISPR technology. **Figure 2C** shows that none of the negative control reactions that were positive using LAMP alone using Eva green were demonstrated to be positive when the LAMP reaction was harnessed with CRISPR.

**Figure 2.**
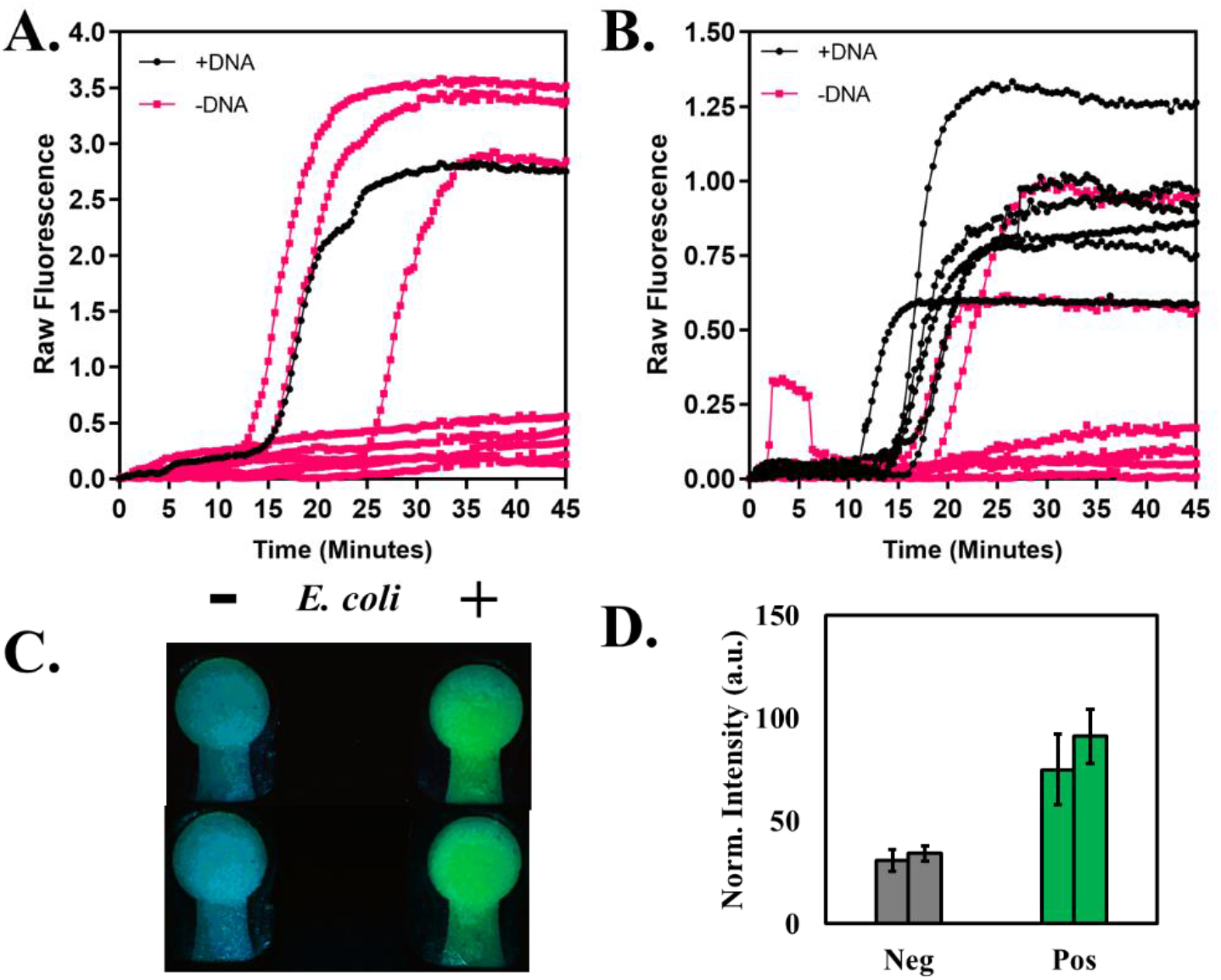
**A**. 3 out of 8 negative control replicates show sporadic nonspecific amplification. **B**. 2 out of 6 replicates of negative controls show non-specific amplification. **C**. Simultaneous CRISPR detection of the non-specifically amplified controls show no green fluorescence on paper confirming that CRISPR specifically differentiates between specific and non-specific amplification. **D**. The normalized intensities of the post amplification image pixels covering the reaction site on paper were analysed to determine mean and standard deviation of the fluorescent signals (N ∼ 1100 pixels) for positive and negative *E. coli* samples.

Microfluidic systems for diagnostics are highly advantageous for their enhanced usability. We therefore moved to verify that the efficiency of CRISPR combined with LAMP in tubes could also be replicated on a chip based fluidic system. We prepared stacks of laser cut Whatman 1 and 113 on PET supporting matrices. The suitability of Whatman 113 over Whatman 1 is demonstrated in **SI Figure S1**.

To incorporate CRISPR-LAMP on paper, after 15 minutes of LAMP at 65°C, the CRISPR cocktail was added onto the paper once the temperature of the reaction block had transitioned to <45°C. Within 5 minutes of adding the CRISPR cocktail, intense fluorescence starts to appear starting at the stalk region of the disc. The paper chip was left to incubate at ∼40°C for the fluorescence signal to noise ratio to noticeably intensify while the reagents diffused uniformly into the paper matrix. Images were captured at 15 minutes (**Figure 3 A**).

**Figure 3.**
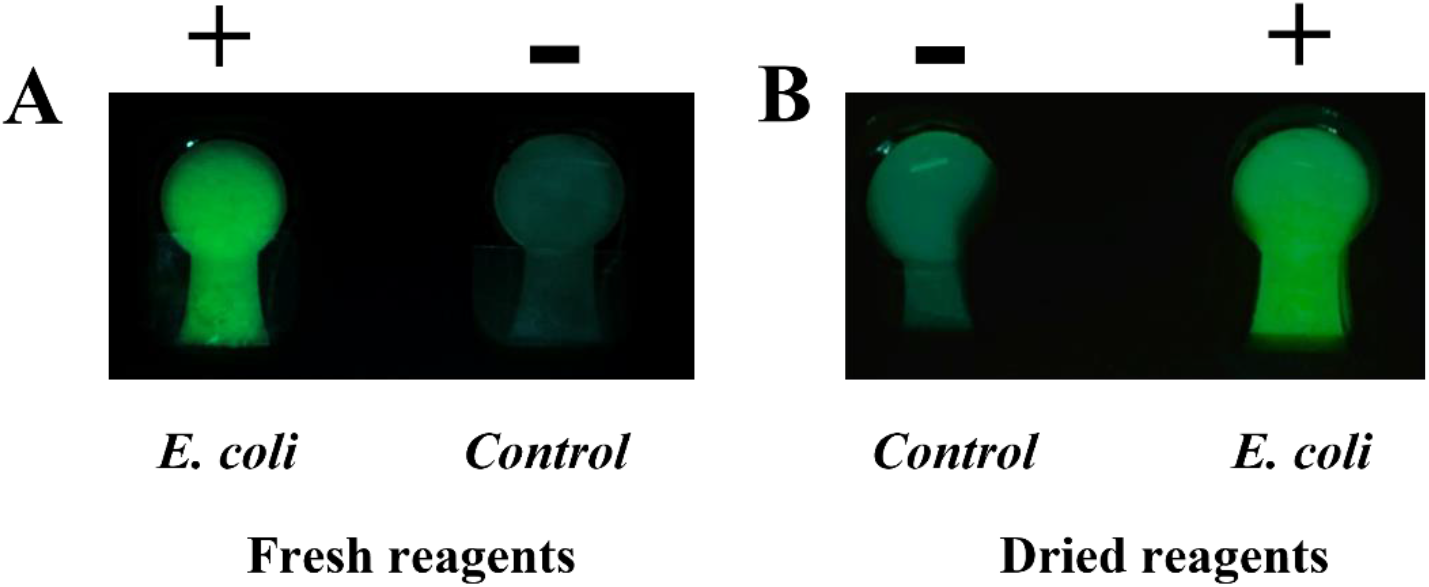
LAMP-CRISPR detection *of E. coli* DNA at ∼20,000 copies/rxn on paper using **A**. fresh reagents and **B**. dried reagents.

The results not only demonstrate the combined power of LAMP-CRISPR on paper, the time to positivity (∼20 minutes) is much reduced when compared to LAMP detection alone (∼40 minutes). Ideally POC testing requires reagents to remain stable without cold chain. Therefore, it was important to test the efficacy of dried LAMP reagents on paper. We found that drying and rehydrating LAMP reagents in the presence of 10% trehalose was suitable for retaining LAMP viability. The CRISPR Cas12 reagents were also dried and rehydrated in tubes to evaluate their viability. As shown in **Figure 3 B**, the CRISPR detection signal is uncompromised when using dried LAMP reagents suggesting amplification efficacy comparable to using fresh reagents. The reaction time with dried reactions was observed to be slightly slower (delayed by ∼5 minutes) compared to using liquid reagents. We speculate that the presence of trehalose in the freeze-dried reagents slows down the diffusion of the reagents.

The successful demonstration of LAMP-CRISPR on a paper substrate shows great potential for the development of NAT technologies for POC settings that are low cost, faster, reliable, and robust.

### Sensitivity studies

To evaluate the linear range and sensitivity of the *E. coli* LAMP assay, analysis was performed with serial dilutions of 40,000 genomic equivalents of E. coli DNA. We found the assay yielded an LOD of 40 copies/rxn tested across replicates in same runs and different runs across different days (**Figure 4 A**).

**Figure 4.**
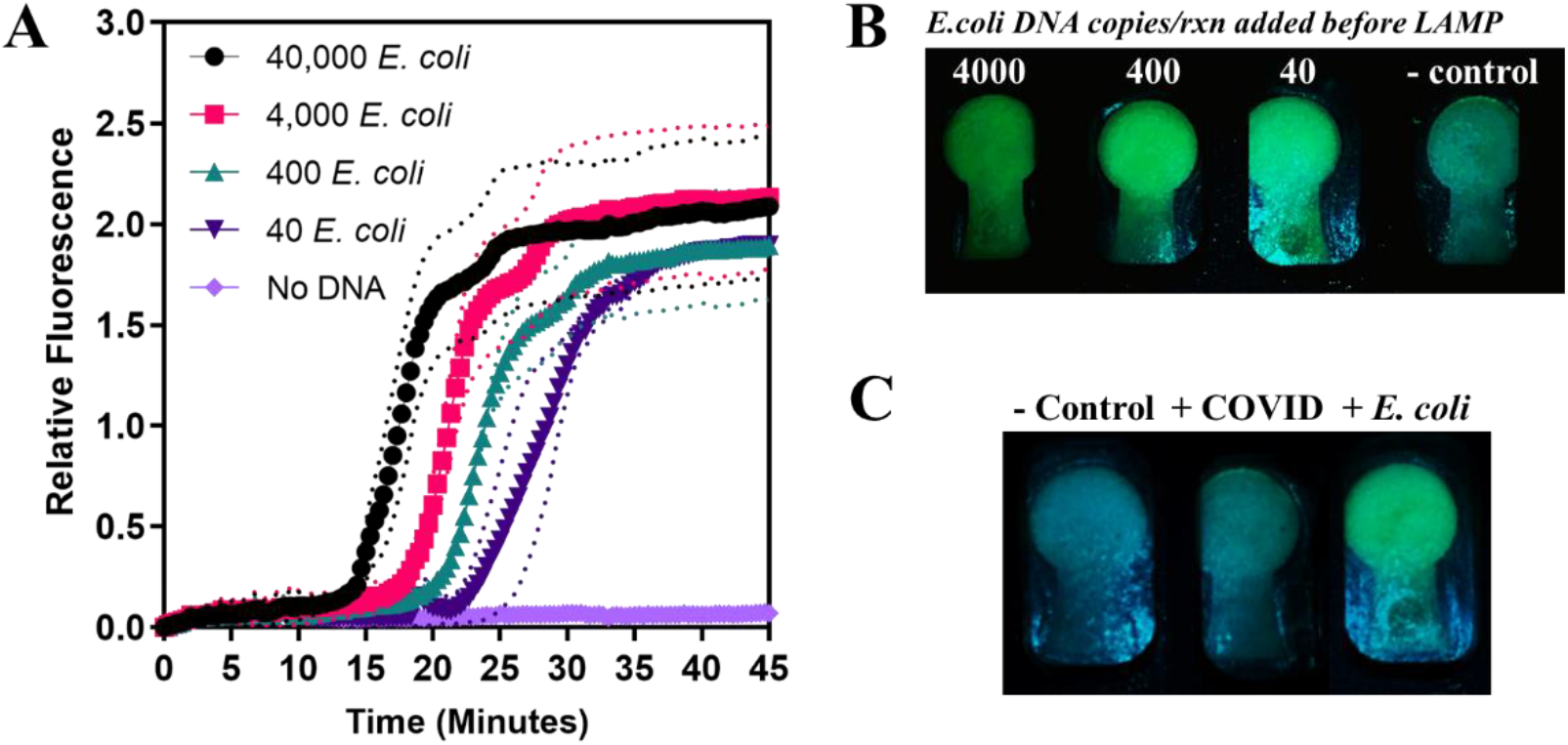
**A**. Raw data of E. Coli LAMP assay that can detect down to 40 copies with an incubation time of 45 minutes. **B**. LAMP on paper amplified 40-4000 copies of target E. Coli DNA and with subsequent CRISPR detection we confirm the successful amplification on paper. LAMP successfully amplified 40 copies of target DNA to the threshold for CRISPR to fire. **C**. Specificity of CRISPR LAMP: LAMP-CRISPR detects E. coli DNA with high specificity while no amplification/detection occurs with COVID DNA and negative control confirming specific detection of *E. coli* DNA.

To evaluate the performance of the *E. coli* LAMP-CRISPR assay on paper, 4000, 400 and 40 copies/rxn were tested. As shown in **Figure 4 B**, the CRISPR detection on paper can successfully detect the amplified product from LAMP starting with as low as ∼40 copies.

### Specificity studies

We evaluated specificity and cross reactivity of the *E. coli* primer set against COVID DNA. LAMP amplification of approximately 10,000 copies/rxn of E.Coli DNA and COVID DNA was detected by CRISPR-Cas12 reaction. As shown in **Figure 4 C**, absence of fluorescence in the COVID head suggested no amplification occurred with COVID DNA during LAMP. Fluorescence observed in the E.Coli head suggested successful amplification reaction.

### POC sample processing and LAMP-CRISPR detection *E*. *coli* in soil

POC diagnostic systems must be robust to handle complex samples. To test the efficacy of our chip system we used soil extracts containing *E. coli* cells. Soil is a complex sample often with an abundance of NAT assay inhibitors. The soil samples were extracted using the S-TECH device as shown in **Figure 5 A** (www.dnaitech.com). **Figure 5 B** shows that *E. coli* DNA extracted from soil samples were successfully detected by the paper chip using LAMP-CRISPR Cas12a detection. The test sample yielded bright fluorescence indicating successful LAMP-CRISPR detection. The control sample (minus E. coli DNA) exhibited no fluorescence. The soil extract E. coli levels were quantified using LAMP in the tube based DNAiTECH Gen2 system (www.dnaitech.com).

**Figure 5.**
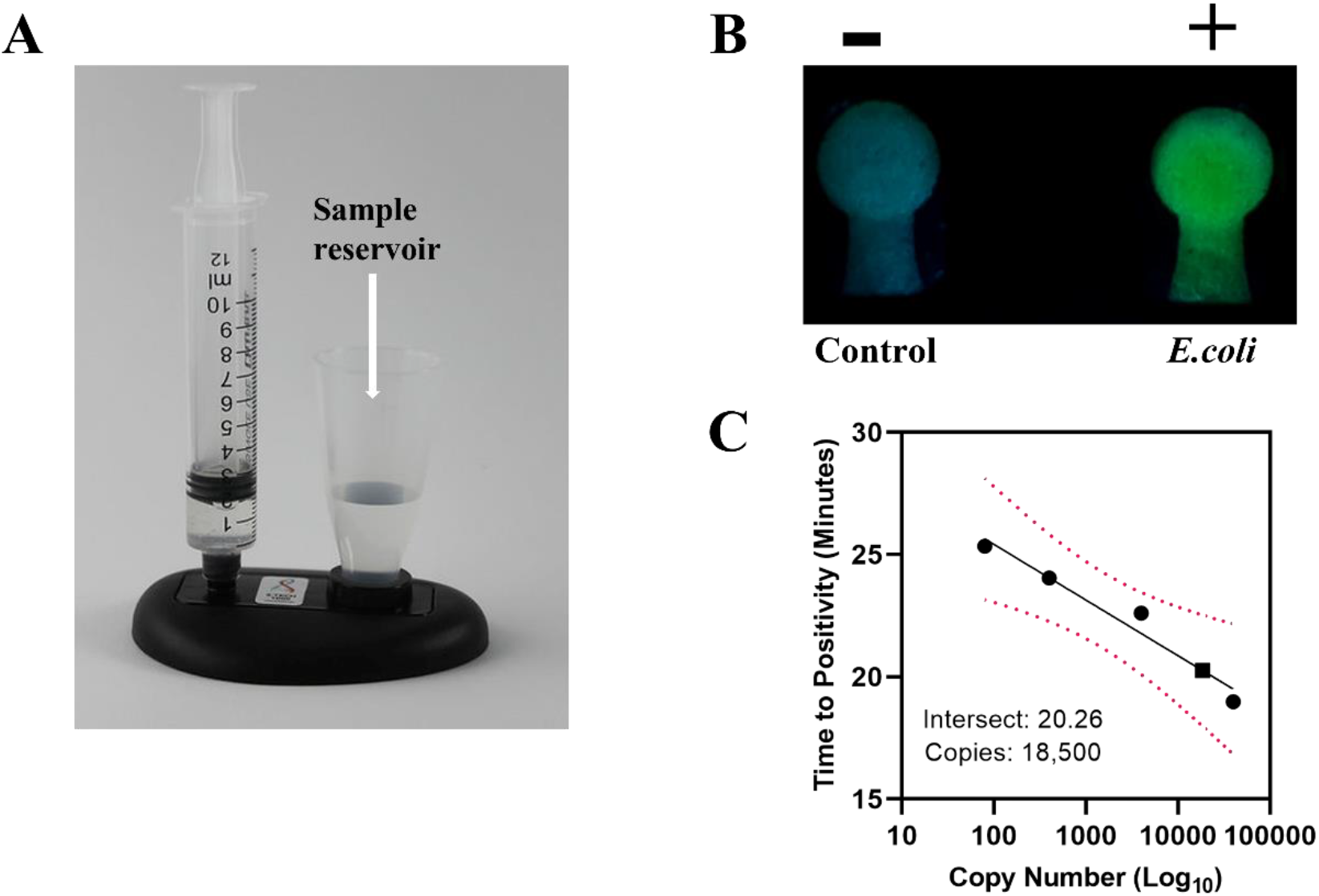
**A**. S-TECH sample preparation device (https://www.dnaitech.com/s-tech) **B**. *E. coli* LAMP-CRISPR fluorescent detection on paper with the soil extract with (right) and without (left) *E. coli* DNA. **C**. Standard curve generated from LAMP assay of *E. coli* DNA recovered using the S-TECH extraction device from soil samples spiked with *E. coli* cells. Black circles indicate the calibration samples of 40,000 GE, 4,000 GE, 400 GE, and 80 GE of *E. coli* genomic DNA. The black square represents the soil sample spiked with *E. coli* cells. The red dots show the 95% confidence interval. The intersect and the copies shown are for the soil extract. The copies shown are per 5μL of soil extract.

The LAMP assay detected ∼18,000 copies/5μL of the extracted DNA sample (**Figure 5C**). The efficiency of the LAMP-CRISPR reaction is evident, the LAMP assay detected presence of *E. coli* DNA in the soil samples in 45 minutes whereas the time to positivity with CRISPR-LAMP detection was only 25 minutes on paper.

This is the first report of a LAMP-CRISPR detection system on paper. Previously all LAMP-CRISPR reactions have been carried out in tubes or in combinations of tubes plus lateral flow devices[8], [30]. Our strategy involves highly specific detection of LAMP amplicons in paper with CRISPR-Cas12a detection system. In our current device, we transition from LAMP to CRISPR by manually adjusting the block temperature from 65 to <45C. Dried CRISPR reagents are then hydrated and flowed into the device at the lower temperature. We envisage future devices that incorporate automatic transition from LAMP reaction temperature to CRISPR reaction temperatures.

## Conclusion

We conclude that the advantage of CRISPR-Cas system combined with existing isothermal amplification technologies can bring NAT technologies to become more efficient, sensitive, specific, and reliable. In this work we achieve LAMP-CRISPR reactions on a paper substrate and provide a first proof of concept paper-based platform for semi quantitative detection of nucleic acids on paper by recording fluorescence signals. We show that LAMP may be prone to non-specific amplification and by using CRISPR we overcome the challenge of false positives with LAMP. We harness the power of CRISPR-Cas 12a to detect LAMP product with high specificity and sensitivity such that with LAMP-CRISPR/Cas12 a system we can detect down to 40 copies of *E. coli* DNA on paper within 30 minutes. With the POC sample processing device (S-TECH), we process soil samples spiked with E. coli cells and report a complete system that is POC deployable. We demonstrate our system is suitable for the detection of various nucleic acids in complex samples. In the future we envision a multiplexed paper-based system that can achieve detection of multiple targets in a single sample.

## Supplementary information

### Testing different paper substrates

We think that Whatman 113 would be a suitable substrate compared to Whatman 1 due to the increased thickness/capacity of Whatman 113 that would result in stronger fluorescent signals. Data shows Whatman 113 to be far superior to Whatman 1 both in terms of reaction viability and visually.

The viability of LAMP assays was also tested by drying and rehydrating LAMP reagents using freeze drying method. We observe that the LAMP enzyme retained activity during drying-rehydration cycles. This was confirmed by successful CRISPR detection of LAMP product in paper.

### LAMP quantification for *E*. *coli* spiked soil sample

**Figure S1.**
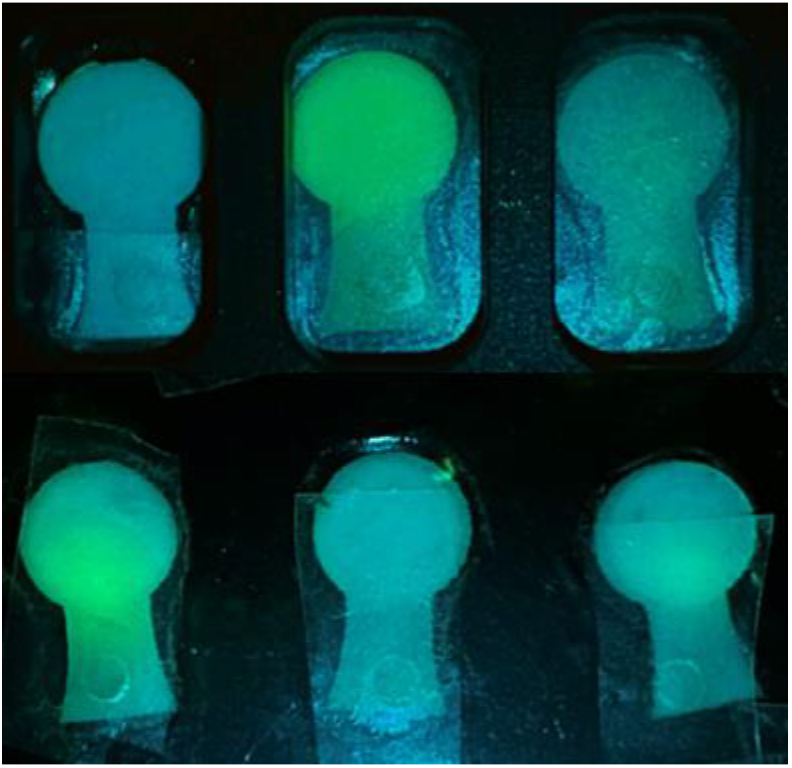
*E. coli* LAMP-CRISPR detection on different paper substrates, Whatman 1 (*right*), 113 (*middle*) and negative control (*left*). Bottom image showing viability of the detection signal when rehydrating dried LAMP reagents on Whatman 1 (*right*) and Whatman 113 (*left*) with negative control (*middle*).

**Figure S2.**
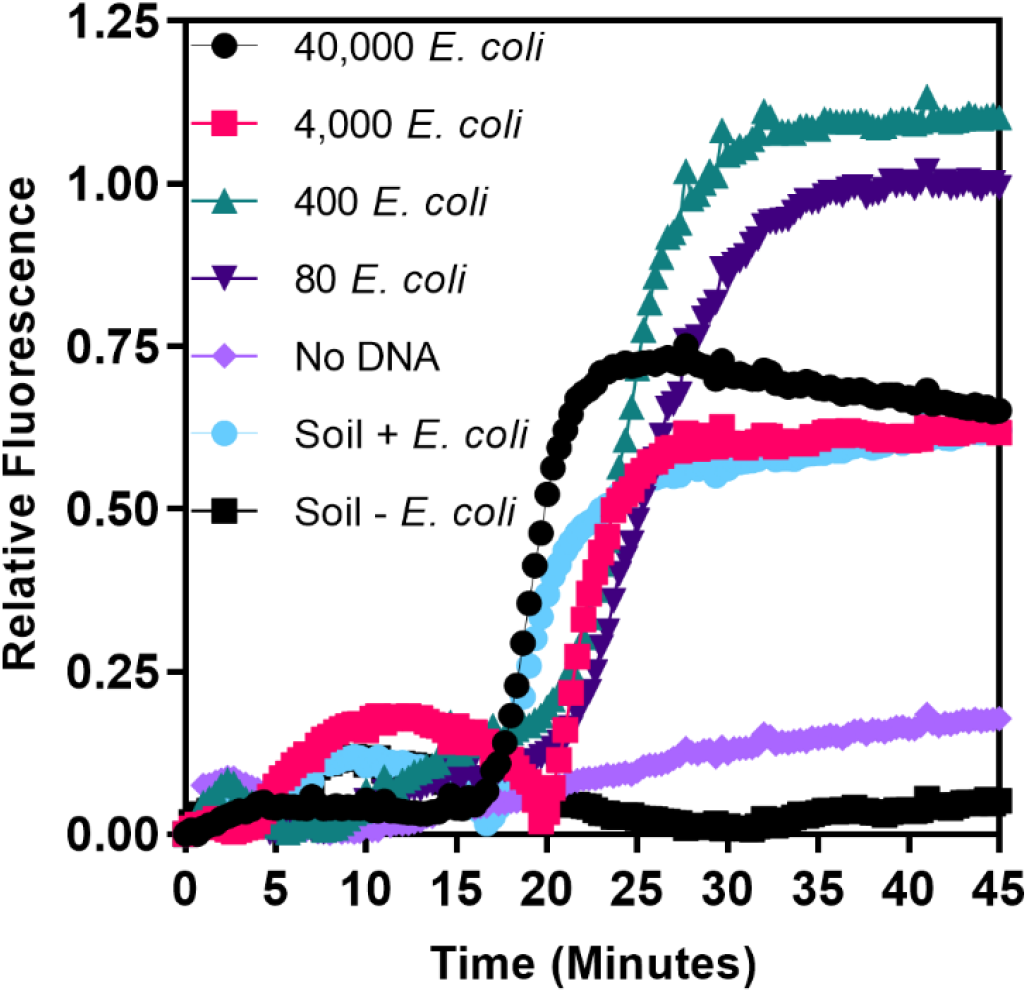
Raw data for the LAMP quantification of *E. coli* spiked soil sample and *E. coli* calibration date for 80-40,000 *E. coli* genomic equivalents.

## Notes

https://www.dnaitech.com/s-tech

https://www.dnaitech.com/dnaitech-gen-2

### Competing Interest Statement

The authors have declared no competing interest.

